# Depleting prion protein using splice-switching small molecules

**DOI:** 10.64898/2026.06.26.734742

**Authors:** Bo Liu, Benjamin Fair, Zhiling Kuang, Zhichao Tang, Junxing Zhao, Lixian Zhou, Qingyao Kong, Aniruddhsingh Solanki, Connor Kenny, James A. Mastrianni, Rui Zhao, Yang I. Li, Jingxin Wang

## Abstract

Prion diseases are fatal neurodegenerative disorders driven by prion protein (PrP) misfolding, and lowering cellular PrP represents a promising therapeutic strategy. Here we report a small-molecule approach that reduces PrP by modulating pre-mRNA splicing of the *PRNP* gene. Through chemical modification of the clinically approved splicing modulator risdiplam, we generated CP3, the first class of compounds that selectively activate a cryptic exon in *PRNP* and routes its mRNA product for degradation, reducing PrP by ∼70% in cells. We demonstrate that CP3 activity critically depends on alternative splicing factor Luc7L, revealing a novel requirement for alternative splicing factors in small molecule splicing modulation. Strikingly, co-administration of CP3 with a Luc7L activator PTC258 significantly lowers PrP levels in the brains of transgenic mice. These results establish splicing modulation as a powerful strategy for PrP reduction and highlight the potential of small-molecule cooperativity for therapeutic RNA targeting.

## Introduction

Prion diseases (PrDs) are fatal neurodegenerative disorders classified as transmissible spongiform encephalopathies.^1,2^ PrD manifests with rapidly progressive dementia, myoclonus, ataxia, and akinetic mutism, leading to death within 5–24 months.^3^ There are currently no approved treatments that modify PrD progression; clinical management remains entirely palliative. PrD is caused by conformational conversion of the cellular prion protein (PrP^C^) into misfolded pathogenic isoforms (PrP^Sc^), which drive templated protein aggregation and neurodegeneration in the central nervous system.^1^

Direct targeting of PrP^Sc^ has been considered impractical due to its aggregation propensity, conformational heterogeneity and ability to circumvent therapeutic intervention by adopting different conformations.^4,5^ In contrast, PrP^C^ has emerged as a rational therapeutic target. Genetic studies demonstrated that conditional ablation of neuronal PrP^C^ in infected mice halts disease progression and mitigates spongiform degeneration, while PrP-null mice are resistant to prion infection without overt neurological or developmental deficits.^67,8^ Human genetic evidence also supports the tolerability of PrP reduction, as large-scale population analyses have identified loss-of-function *PRNP* alleles in adult individuals without evidence of neurological disease,^9^ suggesting that therapeutic reduction of PrP^C^ is safe in humans.

Several therapeutic approaches aimed at reducing PrP^C^ levels have shown promising results, such as anti-PrP^C^ antibodies,^10,11^ which was shown to alter the histopathology in a subset of subjects within a small cohort of PrD patients, suggesting a potential enhancement of prion clearance.^12^ Multiple other strategies, such as antisense oligonucleotides (ASOs),^13,14^ base editing,^15^ and zinc-finger transcriptional repressors^16,17^ have demonstrated disease-modifying effects in preclinical models. Nevertheless, both antibody- and gene-based modalities face challenges in brain delivery and distribution.^18,19^ Small molecules, by contrast, offer distinct advantages in pharmacokinetics.^20,21^ However, PrP^C^ is a classical intrinsically disordered protein (IDP)^22^ and is widely regarded as undruggable by direct protein-targeting approaches.^23^ Targeting PrP expression at the mRNA level offers a structure-independent approach that can exploit RNA splicing modulation to reduce cellular PrP levels.

Like many human genes, *PRNP* harbors a long non-coding intron (12.7 kb) that must be precisely removed during pre-mRNA splicing to generate a functional transcript. We hypothesized that a splice-switching small molecule could disrupt this step and thereby deplete the productive *PRNP* mRNA. After screening a panel of reported splicing modulators, we found that risdiplam, an FDA-approved compound that selectively engages the 5′ splice site (ss) of the *SMN2* gene (unrelated to PrD), can alter the splice pattern of *PRNP* at a high concentration.

Structural and biochemical analyses indicate that risdiplam functions as a molecular glue that binds at the interface between pre-mRNA and a spliceosome sub-complex U1 small nuclear ribonucleoprotein (U1 snRNP), stabilizing this ternary complex. Risdiplam enhances recognition of weak 5′ ss characterized by a –2G –1A motif immediately upstream of the exon–intron junction, commonly referred as the GA|GU 5′ ss (“|” denotes splice junction) (Extended Data Fig. 1).^24,25^ This sequence motif is shared by multiple splicing modulator-responsive genes, including *SMN2*, *HTT*, and *FOXM1*.^20,26–28^ In contrast, our analysis revealed that at high concentrations, risdiplam engages a distinct 5′ ss within *PRNP* with the GU|GU motif (Extended Data Fig. 2), a significantly weaker target identified from previous reports.^25,29^

Through structure-activity-relationship studies, we developed CP3, a compound derived from the risdiplam scaffold, with an altered 5′ ss recognition preference that effectively targets the GU|GU motif. In human glioblastoma cells, CP3 selectively promotes inclusion of a *PRNP* cryptic exon, resulting in a reduction of PrP^C^ protein levels through nonsense mediated decay (NMD). Furthermore, we discovered that kinetin analogs can synergize with CP3 by upregulating Luc7L, an alternative splicing factor essential for CP3 function. Co-treatment of CP3 and kinetin analogs such as PTC258 increased the potency of CP3 by ∼10-fold, substantially lowering the drug concentration needed to achieve clinically relevant PrP suppression. Furthermore, co-administration of CP3 and kinetin analogs significantly changed *PRNP* splice pattern in transgenic mice expressing human *PRNP*, leading to significant PrP suppression. Together, these findings establish a strategy for selective PrP suppression through splicing modulation and highlight small-molecule cooperativity as a broadly applicable strategy for overcoming undruggable protein targets.

## Result

### Reprogramming risdiplam to target a cryptic GU|GU 5′ ss in *PRNP*

To shift the risdiplam scaffold toward selectively targeting of GU|GU 5’ ss, we screened a collection of 126 structurally diverse risdiplam analogs using a GU|GU 5′ ss reporter gene assay and compared their activities with GA|GU and other 5’ ss variants. We observed that GU|GU 5′ ss responded to a significant subset of risdiplam analogs (Fig. 1A) and identified the most selective GU|GU hit in the 126-compound library, namely SM74. NMR studies further suggested that risdiplam analogs use the middle two rings (denoted as the core) to interact with the bulged A.^25^ Our previous molecular dynamics and NMR studies also demonstrated that the core of risdiplam scaffold is critical for mediating hydrogen-bond interactions with other types of bulged nucleotides in duplexed RNAs.^30^ We therefore hypothesize that the core of risdiplam analogs determines 5′ ss selectivity, and additional substituents on the central bicyclic core compensate for the size difference between A and U. This hypothesis is supported by our GU|GU-selective screening hit SM74, in which a bulky, cyclic substituent on the core structure markedly shifts selectivity toward GU|GU over GA|GU 5′ ss (Fig. 1B, 1C).

**Fig. 1:**
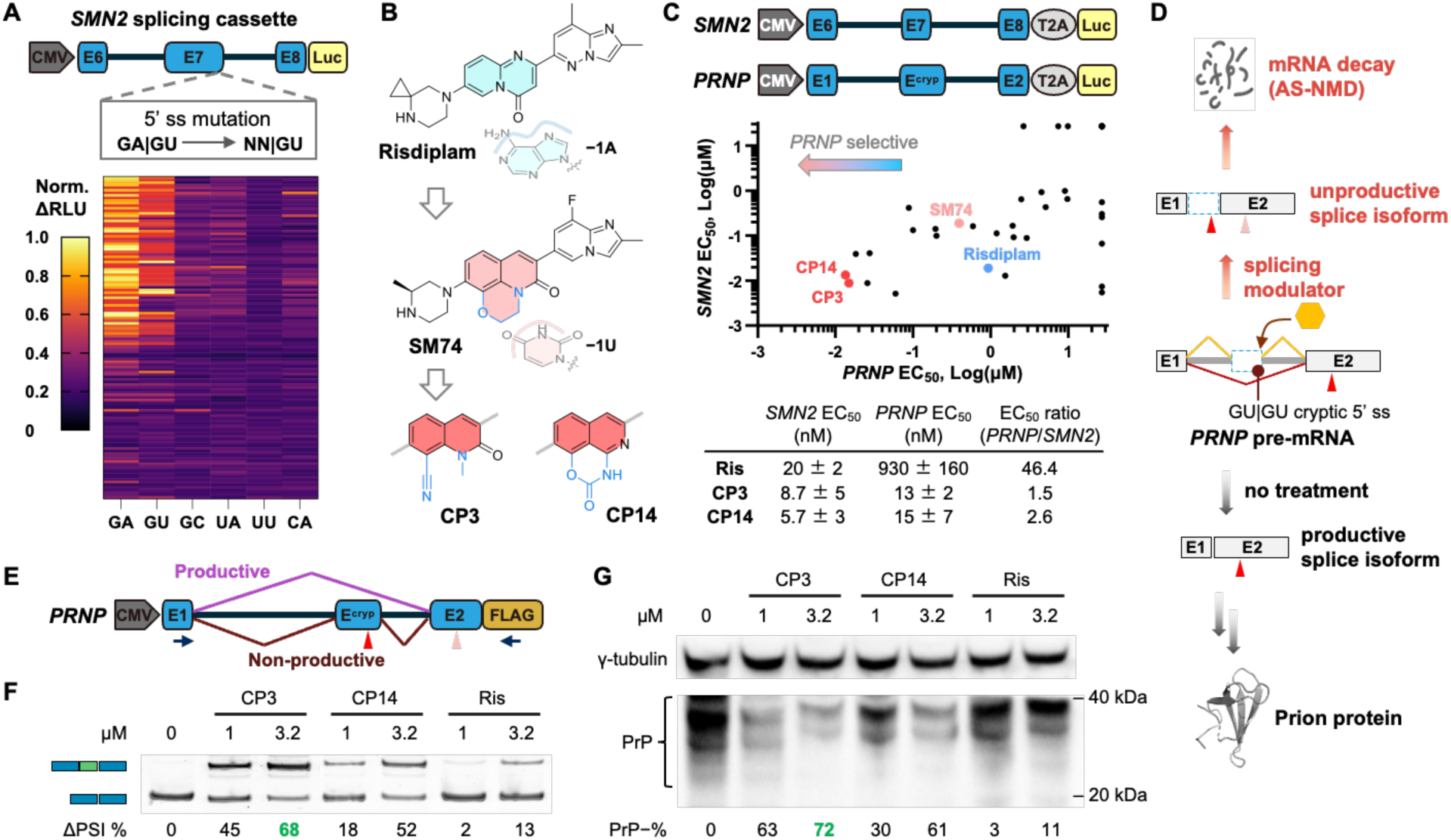
Expanding target space of splicing modulation enables a new approach for the treatment of prion disease. (A) Design of splicing minigene assays for 5′ ss variants (NN|GU) in the *SMN2* splicing cassette (Luc, Gaussia luciferase). Heatmap representing the normalized relative light unit signal change (norm. ΔRLU) of the assay at 5 µM for 126 risdiplam analogs (36 h, 293T cells). (B) Structures of risdiplam, SM74, CP3, and CP14. Modifications to the central bicyclic rings likely accommodate the size difference between A and U bulges. (C) Reporter minigene assay constructs for *PRNP* (GU|GU) and *SMN2* (GA|GU) splicing. Scatter plot illustrating the splicing modulatory activities of 38 analogs. The table shows EC_50_ values and relative selectivity for risdiplam, CP3, and CP14 determined in these assays. (D) Biological outcomes of poison cryptic exon inclusion in the *PRNP* gene induced by splicing modulation, leading to *PRNP* transcript degradation via the AS-NMD pathway (red arrow, stop codon). (E) *PRNP* minigene construct used to measure dose-response effects of splicing modulators. (F) *PRNP* splicing changes assessed by end-point RT-PCR, and visualized and quantified by PAGE densitometry (ΔPSI, percent spliced in difference). Representative gel image of three independent experiments. (G) PrP expression suppression measured by anti-PrP Western blot (multiple bands, glycosylation states; PrP−%, percent reduction in PrP levels, quantification by densitometry) (24 h, 293T cells). All data are representative of three independent experiments.

RNA-seq experiments demonstrated that SM74 can induce the inclusion of a cryptic “poison” exon situated within intron 1 of the *PRNP* gene, generating a non-productive splice isoform with a premature stop codon (Fig. 1D). This cryptic exon inclusion activates the alternative splicing-coupled NMD (AS-NMD) pathway, thereby reducing the overall *PRNP* mRNA level. We synthesized 38 analogs of SM74 to further probe the core structure and screen for compounds with enhanced potency. To evaluate the splicing activity of these analogs, we constructed two reporter gene assays for *SMN2* and *PRNP* pre-mRNA splicing with GA|GU and GU|GU 5′ ss, respectively (Fig. 1C). Compound-induced exon inclusion drives the luciferase out of frame, resulting in a reduction of luciferase signal. By plotting the half-maximal effective concentration (EC_50_) from the two assays, we identified CP3 and CP14 (Fig. 1C) for their superior potency and selectivity towards *PRNP* over *SMN2* (Fig. 1C). In this set of minigene splicing assays, both compounds are >60 times more potent than risdiplam in modulating *PRNP* splicing while maintaining similar activity on *SMN2* (Fig. 1C). Consistent with SM74, both CP3 and CP14 contain substituents on the core scaffold with stronger electron-withdrawing properties (Fig. 1B).

To further assess the impact of these compounds on splicing pattern changes and protein levels, we used a *PRNP* splicing vector containing the full PrP coding sequence fused with a FLAG tag (Fig. 1E). In this system, CP3 demonstrated a 68% change in the *PRNP* splice pattern (Fig. 1F) and a 72% reduction in PrP levels (Fig. 1G, Extended Data Fig. 3). We transfected the *PRNP* splicing vector into mouse Neuro-2A cells and observed similar potency in PrP reduction (Extended Data Fig. 4). Because CP3 performed slightly better than CP14 in PrP-lowering activity with a lower cytotoxicity (Fig. 1F, 1G, Extended Data Fig. 5), we used CP3 as the lead compound in the following experiments.

### Validation of potency and selectivity of CP3 in wild-type cells

We first validated CP3 activity on endogenous PrP protein in a human glioblastoma cell line LN229, which exhibits relatively high levels of PrP expression.^31^ Treatment of LN229 cells with CP3 resulted in a maximal ∼70% reduction in *PRNP* mRNA levels, consistent with the observed splicing changes (Fig. 2A, 2B). Dose-response analysis using Western blot also demonstrated that 3.2 μM CP3 reached a plateaued effect to reduce PrP level by ∼70 % in 24 h (Fig. 2C).

**Fig. 2:**
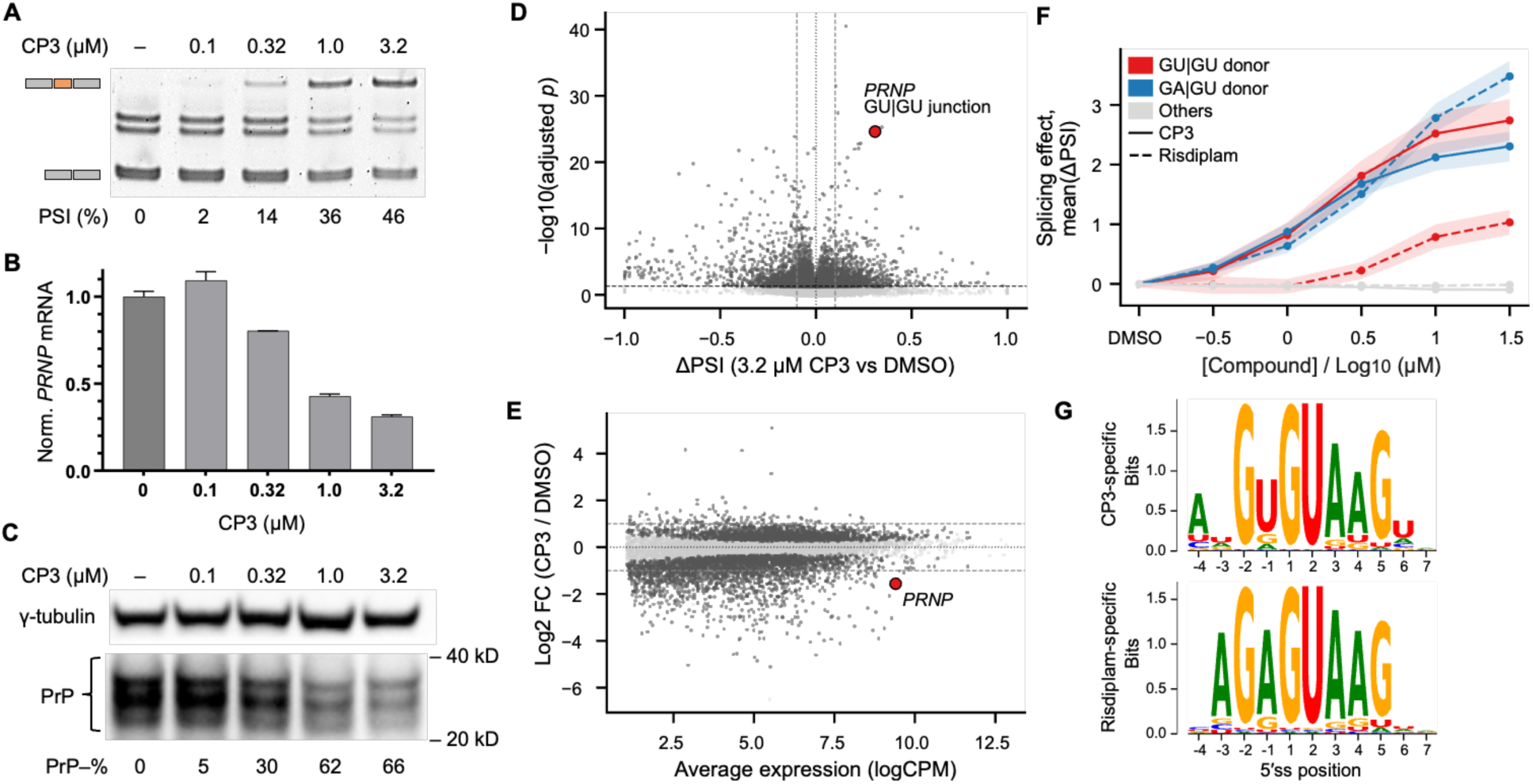
CP3 lowered PrP expression in wild-type human glioblastoma cells (LN229). (A) *PRNP* splicing patterns in cells treated with CP3 (24 h) were analyzed by end-point PCR. PCR products corresponding to productive and nonproductive splice isoforms were quantified by PAGE densitometry (PSI, percent spliced in difference). (B) Quantification of *PRNP* mRNA level in cells treated with CP3 by RT-qPCR (24 h). (C) Anti-PrP Western blot analysis of PrP^C^ levels following dose-response treatment with CP3 (36 h). PrP levels were normalized to γ-tubulin levels, and percentage reduction (PrP–%) was quantified by PAGE densitometry. (D) Differential splicing volcano plot (3.2 µM CP3 vs DMSO). Dashed lines indicate adjusted P = 0.01 and ΔPSI = ±10%. The *PRNP* GU|GU cryptic junction is highlighted in red. (E) MA plot of differential gene expression (3.2 µM CP3 vs DMSO). Genes with FDR < 0.01 are highlighted in red, including the labeled *PRNP* gene. (F) Meta-splice site dose-response plots showing ΔPSI relative to DMSO control, averaged across leafcutter-detected splice donor sites (GA|GU: n=5,560; GU|GU: n=3,261; others: n=180,894). (G) Seqlogos of 5′ ss from CP3-specific (*n* = 266), and risdiplam-specific (*n* = 69) events, defined by differences in EC_50_ between risdiplam and CP3 (local false sign rate < 0.01) and filtered for −2G to enrich for direct splicing modulation effects.

We next assessed the selectivity of CP3. We treated LN229 cells with CP3 at a series of concentrations and performed RNA-seq. Splicing analysis using LeafCutter^32^ revealed the *PRNP* GU|GU 5′ ss junction as one of the most significantly activated splicing events in 3.2 μM CP3-treated cells (Fig. 2D). When comparing gene expression changes at a dose that achieves ∼2-fold downregulation, *PRNP* exhibited the largest reduction among genes expressed at comparable or higher levels, indicating drug sensitivity (Fig. 2E).

We further characterized the global impact of CP3 on RNA splicing. Meta-analysis of splice donor sites detected revealed that GU|GU sites, as a class, are activated to a similar extent as GA|GU sites in a concentration-dependent manner by CP3, whereas other splice sites are, on average, unaffected (Fig. 2F). We then fit dose-response curves to estimate EC_50_ values for individual splice junctions. Consistent with our meta-splice site dose response curves, we found that the risdiplam and CP3 exhibit similar EC_50_ values for GA|GU splice sites, whereas for GU|GU splice sites, risdiplam shows an ∼16-fold higher EC_50_ than CP3 (Extended Data Fig. 6). To elucidate the consensus 5′ ss activation motif of CP3, we aggregated all CP3-activated splice sites with EC_50_ < 3.2 μM and compared them with those for risdiplam under a −2G constraint. The activation motif for CP3-activated splicing events is ANGW|GTAAG (W = U/A) (Extended Data Fig. 7). In contrast, risdiplam-specific splicing more exclusively favors –1A in its activation motif, consistent with prior reports (Extended Data Fig. 7).^29^ For CP3-specific splice junctions (EC_50_[CP3] < EC_50_[risdiplam] with statistical significance), the activation motif reveals selective activation of −1U, as well as a stronger specificity for −4A (Fig. 2G). Collectively, these findings indicate CP3 is a selective PrP-lowering candidate targeting GU|GU 5′ss in human cells.

### CP3 activity is enhanced by kinetin analogs

To further improve CP3 activity, we screened a panel of known splicing modulators in the presence of CP3 at a low concentration (0.32 µM). Notably, we discovered that orally available, brain penetrant kinetin analogs such as RECTAS^33,34^, BPN-15477^35^, and PTC258^36^ can significantly improve the activity of CP3 in cells. BPN-15477 (25 μM) or PTC258 (32 nM) can potentiate CP3 at 0.32 μM to achieve the splice pattern change similar to that observed with CP3 at 3.2 μM (Fig. 3A). Accordingly, the combination of BPN-15477 or PTC258 can also robustly reduce PrP at both the mRNA (Fig. 3B) and protein levels (Fig. 3C) with 0.32 μM CP3, decreasing the CP3 dose required for maximum PrP reduction by ∼5-fold. On the other hand, BPN-15477 or PTC258 alone has minimal effects on the *PRNP* gene in human cells (Extended Data Fig. 8A). Volcano plot of splice pattern changes in LN229 cells treated with a combination of CP3 and BPN-15477 confirmed that *PRNP* remains one of the most responsive genes to this treatment (Extended Data Fig. 8B). Genome-wide RNA-seq analysis suggests a global potentiation effect of BPN-15477 on CP3, with an average 7.7-fold increase in potency upon combination treatment (Extended Data Fig. 8C). The presence of these kinetin analogs at the effect concentration did not significantly affect the cytotoxic profile of CP3 in LN229 cells, suggesting that this is a promising combination approach to enhance the PrP reduction in cells and in vivo.

**Fig. 3:**
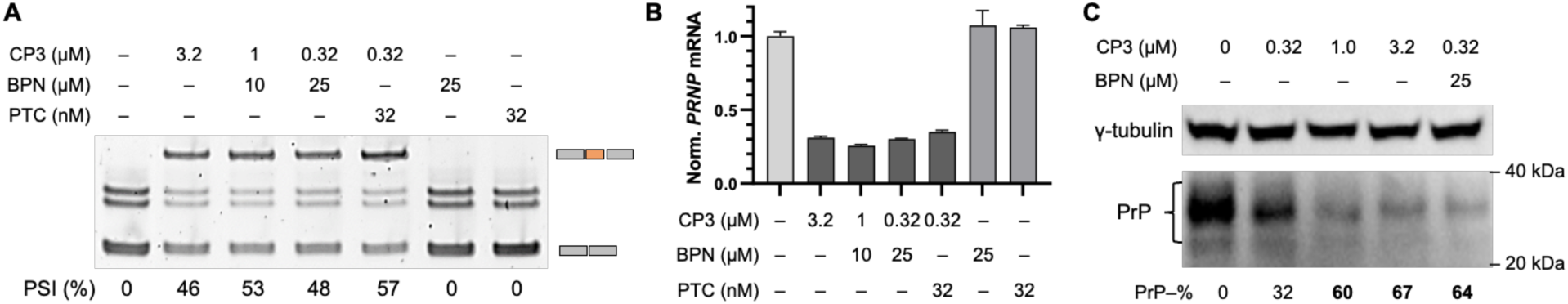
Kinetin analogs potentiate CP3 activity in LN229 cells. (A) *PRNP* splice patterns in cells treated with CP3 in combination with kinetin analogs at various concentrations, assessed by end-point RT–PCR (24 h). (B) Quantification of *PRNP* mRNA reduction in cells treated with splicing modulator combinations by RT–qPCR (24 h). BPN, BPN-15477; PTC, PTC258; error bars, s.d. (n = 3). (C) Anti-PrP Western blot analysis of LN229 cells treated with BPN-15477 in combination with CP3 at various concentrations (48 h). Data are representative of three independent experiments.

### Luc7L is essential for the effect of CP3 on 5′ ss recognition

We next sought to investigate the mechanism by which these kinetin analogs synergize with the activity of CP3. BPN-15477 was shown in a report to activate CLK1 and upregulate SRSF6 phosphorylation to enhance RNA splicing.^34^ The mechanism of action of PTC258 remains to be elucidated. Cells treated with BPN-15477or PTC258 showed an upregulation of alternative splicing factor Luc7L by ∼4-fold (Extended Data Fig. 9A), consistent with reported datasets on kinetin analogs.^35,37^ Proteins in the Luc7L family (Luc7L/L2/L3) are homologs of the yeast Luc7 protein, which binds the U1 snRNA–5′ ss RNA duplex opposite U1C and stabilizes the duplex, together with U1C^38^. Recent studies show that Luc7L family proteins regulate 5′ ss selection in a sequence-dependent manner, with Luc7L/L2 promoting sites with stronger intronic consensus and Luc7L3 promoting sites with stronger exonic consensus.^39^ Our splicing analysis of BPN-15477-treated cells revealed that the *LUC7L* gene contains a naturally occurring poison exon within intron 1, leading to a nonproductive transcript. BPN-15477 treatment corrected this mis-splicing event, thereby increasing Luc7L expression (Extended Data Fig. 9B).

To test whether Luc7L improves CP3’s activity on *PRNP* 5′ ss, we reconstituted human U1 snRNP using recombinant proteins and in vitro transcribed U1 snRNA^40^. Using a fluorescence polarization assay with a fluorophore-labeled RNA containing the *PRNP* GU|GU 5′ ss, we found that reconstituted U1 snRNP binds relatively weakly to the *PRNP* 5′ ss (Fig. 4A). The presence of CP3 does not improve the binding affinity between reconstituted U1 snRNP and the *PRNP* 5′ ss (Fig. 4A), consistent with our previous finding that risdiplam does not affect the binding affinity between reconstituted U1 snRNP and the *SMN2* 5′ ss while branaplam (a different class of splicing modulator) does^40^. These findings indicate that an additional cellular factor outside the reconstitution system is required for the activity of risdiplam and CP3. We then expressed and purified human Luc7L, L2, and L3 without their RS domain in *E. coli*. We demonstrated that in the presence of Luc7L or L2, but not L3, CP3 improves the binding between reconstituted U1 snRNP and the *PRNP* 5′ ss (Fig. 4A, 4B). When the weak GU|GU 5′ ss in *PRNP* pre-mRNA is mutated to a strong site (GG|GU), the binding affinity between the mutant 5′ ss and U1 snRNP is markedly enhanced but indiscriminative between CP3-treated samples and the negative control even in the presence of Luc7L/L2 (Fig. 4A), indicating that compound activity is specific to the GU|GU 5′ ss. Similarly, Luc7L/L2 is required for risdiplam to enhance the interaction between reconstituted U1 snRNP and the *SMN2* 5′ ss (Extended Data Fig. 10), indicating a broadly applicable mechanistic requirement for risdiplam-like molecules.

**Fig. 4:**
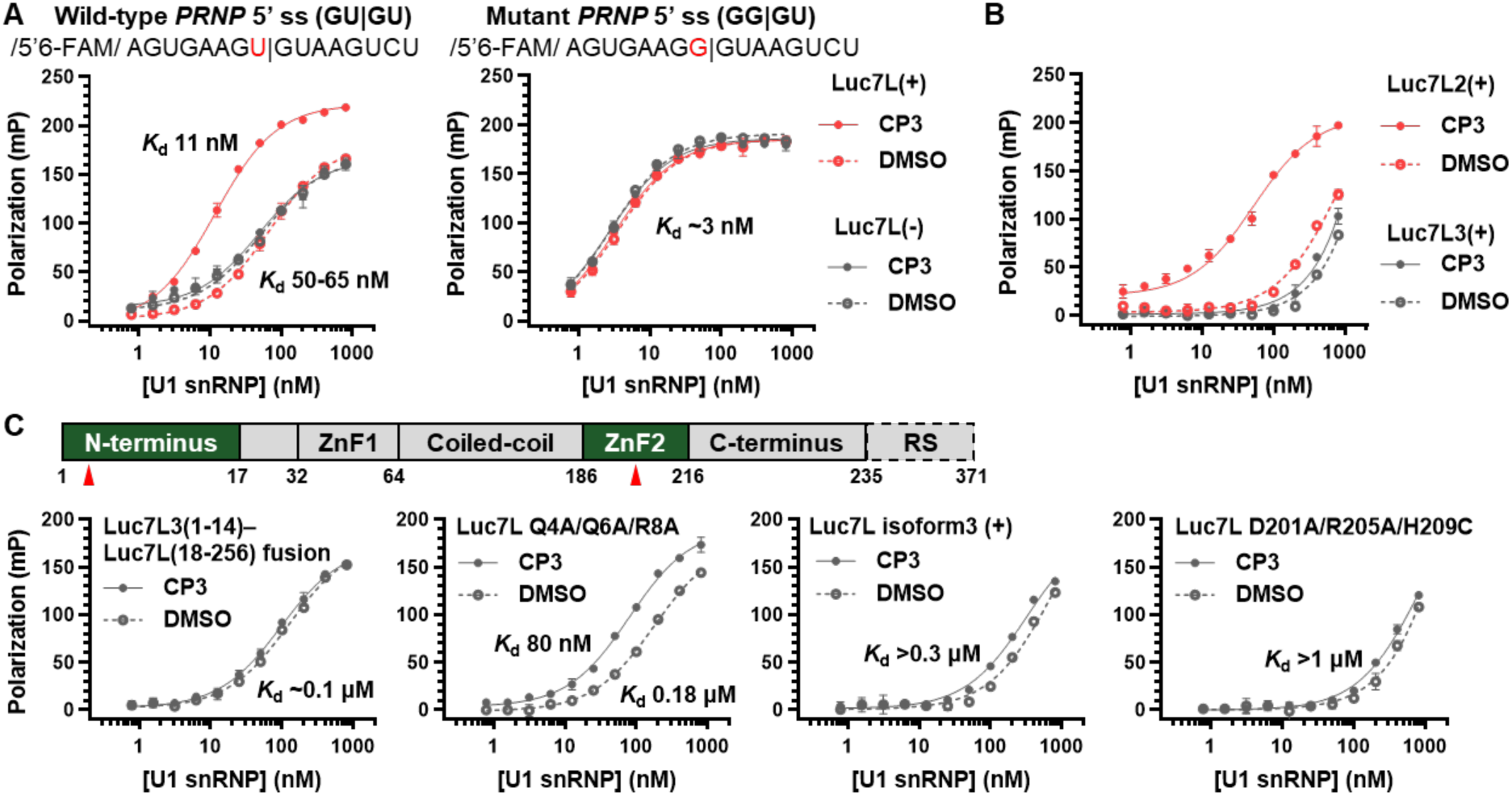
Luc7L or Luc7L2 is a required component for U1 snRNP−*PRNP* 5′ss−CP3 complex formation. In vitro binding of U1 snRNP to *PRNP* pre-mRNA oligonucleotide (5’ 6-FAM-tagged) containing 5′ss in the presence of CP3 and **(A)** with or without Luc7L (10 μM) in fluorescence polarization (FP) assays. A mutant *PRNP* pre-mRNA oligonucleotide with GG|GU 5′ss is included for negative control. Estimated *K*_d_ values from curve-fitting are shown beside each curve, and **(B)** with Luc7L2 or Luc7L3 (10 μM) in fluorescence polarization assays. Estimated *K*_d_ values from curve-fitting: Luc7L2(+)–CP3, 53 nM; Luc7L2(+)–DMSO, >0.5 μM; Luc7L3 curves, >1 μM. **(C)** Domain architecture of Luc7L, highlighting key residues required for CP3 function by domain swapping and mutagenesis experiments. In vitro binding experiments were performed using FP assays in the presence of various Luc7L variants in *N*-terminus and ZnF2 domains (see A–B for conditions). Red arrow, mutagenesis locations. Estimated *K*_d_ values from curve-fitting are shown beside each curve. Data are shown as mean ± s.d. from three technical replicates.

We next sought to understand the reason why Luc7L/L2 but not L3 is crucial for CP3 binding. We performed multiple sequence alignment of the three proteins, which shows that Luc7L and Luc7L2 are more similar to each other than to L3 in the non-RS region (Extended Data Fig. 11). Using domain swapping experiment, we found that replacing the *N*-terminal 14 amino acids of Luc7L that are predicted to be a helix^41^ with those of Luc7L3 abolishes the effect of Luc7L on CP3 (Fig. 4C). Interestingly, Luc7L has an isoform 3 that lacks this *N*-terminal helix (Extended Data Fig. 11) and this isoform does not enable CP3 to improve the binding between U1 snRNP and *PRNP* 5′ ss (Fig. 4C), consistent with the importance of the *N*-terminal helix of Luc7L in its effect on CP3. Multiple sequence alignment revealed substantial differences between the *N*-terminal region of Luc7L/L2 and L3 (Extended Data Fig. 11). We generated a fusion protein containing the Luc7L3 N-terminus and the remainder of Luc7L, as well as a Luc7L triple mutant (Q4A/Q6A/R8A) and found that both variants markedly reduced Luc7L-dependent CP3 activity (Fig. 4C), supporting the importance of the *N*-terminal helix in its effect on CP3. An AlphaFold3 prediction indicates that the *N*-terminal helix interacts extensively with the Sm ring (Extended Data Fig. 12), potentially help anchoring Luc7L to U1 snRNP.

In addition, the AlphaFold3 prediction indicates that the ZnF2 domain of Luc7L interacts with the U1 snRNA and 5′ ss duplex, mimicking the interaction between yeast Luc7 ZnF2 and the RNA duplex^38^. This prediction suggests that residues on Luc7L ZnF2 are potentially also important for its effect on CP3 in human cells. To test this hypothesis, we expressed and purified a Luc7L triple mutant D201A/R205A/H209C corresponding to residues on yeast Luc7 that interact with the RNA duplex.^38^ As expected, this Luc7L triple mutant abolished the effect of Luc7L on CP3 (Fig. 4C), supporting our hypothesis.

### CP3–PTC258 combination reduced PrP expression in vivo

To validate the in vivo activity of CP3, we first evaluated its effects in primary neuronal cultures derived from neonatal transgenic mice (Tg26372), which harbor 20 tandem copies of the human *PRNP* transgene on a *Prnp*^−/−^ background.^42^ Treatment with CP3 for 24 hours resulted in a shift in *PRNP* splice isoform distribution toward the nonproductive transcript, with a reduction in PrP protein levels (Fig. 5A, 5B). To extend these findings in vivo, CP3 was administered by intracerebral injection to neonatal mice. We selected PTC258 as an adjuvant based on its superior ability to potentiate CP3 activity in cell culture (Fig. 3A). Analysis of brain tissue demonstrated a splice-switching effect comparable to that observed in primary neuronal cultures (Fig. 5C). Encouraged by the in vivo splicing modulation observed following drug treatment, we next examined prion protein expression in mice receiving two intracerebral injections of CP3/PTC258 combination. As shown in Fig. 5D, Western blot analysis revealed a significant reduction in PrP levels compared with the vehicle-treated group. Quantitative analysis demonstrated an ∼40% reduction in prion protein expression in treated mice (Fig. 5E), with the potential for greater reduction over longer treatment durations^43^ and conferring phenotypic effects.^13^

**Fig. 5:**
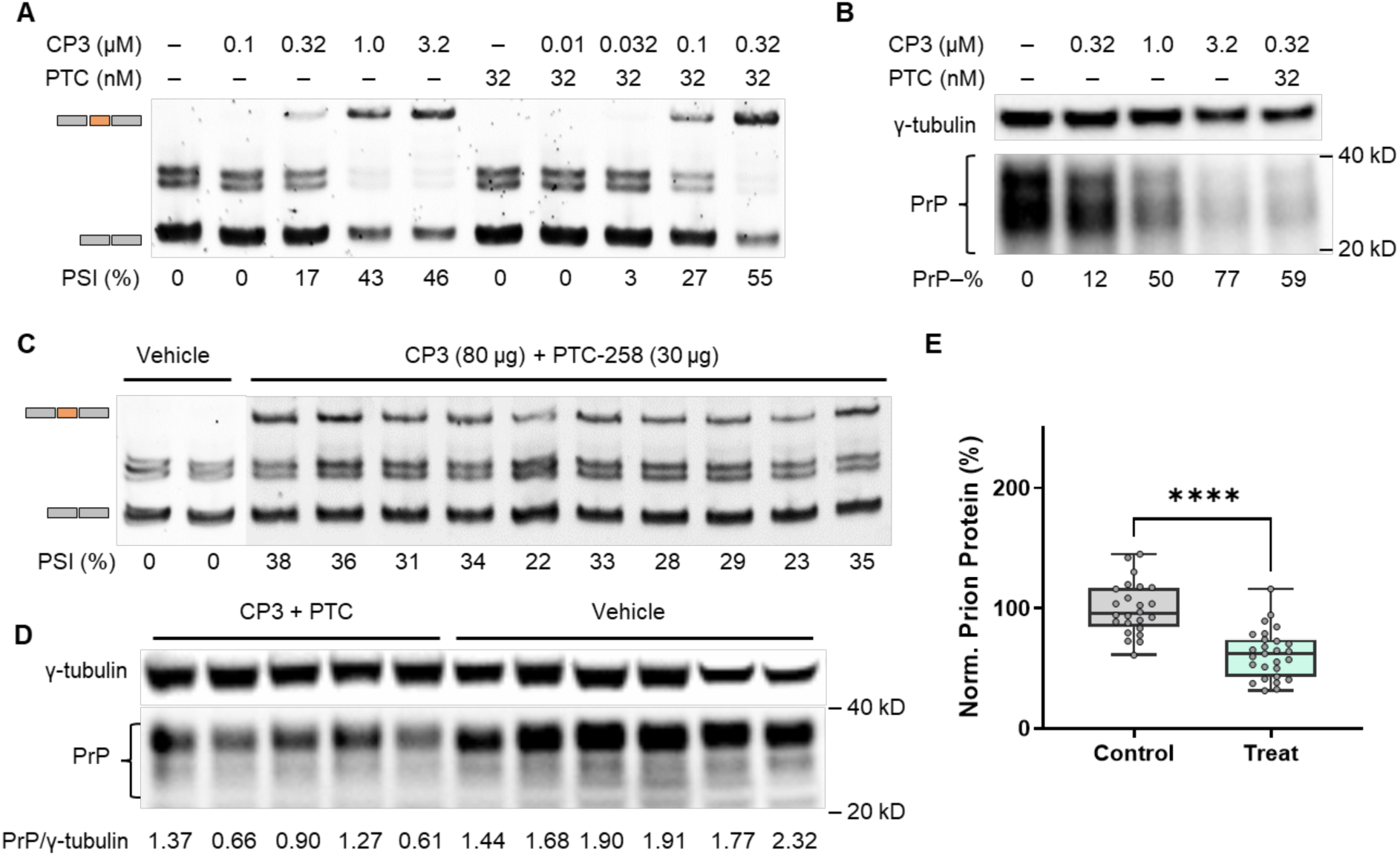
Activity of CP3 and its combination with PTC258 in human PrP transgenic mice (Tg26372). (A) Endpoint RT-PCR analysis of *PRNP* splicing in primary neuronal cultures (P1–3) derived from Tg26372 mice following 24 h treatment with various concentrations of CP3, alone or in combination with PTC258 (32 nM). (B) Anti-PrP Western blot analysis of the same samples as in (A). (C) Endpoint RT-PCR analysis of *PRNP* splicing in brain tissues of Tg26372 mice (P1–3) following single intracerebral administration of CP3 (80 μg) in combination with PTC258 (30 μg) for 24 h (vehicle *n* = 2; compound treatment *n* = 10). (D) Anti-PrP Western blot analysis of brain tissues from Tg26372 mice (P1–P3) after intracerebral administration of CP3 (80 μg) and PTC258 (30 μg) for 48 h (2 doses, 1 dose/day). Representative samples are shown out of 24−27 mice per group. (E) Quantification of normalized prion protein levels (Western blot) in control and treated mice (*n* = 24−27 per group). Each dot represents an individual sample. Boxes, median and interquartile range. Whiskers, range of data. ****p < 0.0001 (unpaired two-tailed test). For (A) and (C), PCR products corresponding to productive and nonproductive splice isoforms were visualized and quantified by PAGE densitometry (ΔPSI, percent spliced in difference). For (B) and (D), PrP levels were normalized to γ-tubulin levels, and percentage reduction (PrP–%) was quantified by PAGE densitometry.

## Discussion and Conclusion

Together, our results establish a first-in-class compound that drives splicing-dependent PrP reduction both in vitro and in vivo, providing a new therapeutic strategy for prion diseases. The lead compound CP3 selectively engages a GU|GU 5′ ss, reduces *PRNP* mRNA and protein levels by ∼70% in human cells, and exhibits favorable pharmacodynamic properties in vivo. Strikingly, the combination of CP3 with kinetin analogs enhances activity and reduces the required dose, thereby minimizing potential off-target toxicity.

Our data demonstrate the exciting possibility of reprogramming the specificity of small molecule splicing modulators for the treatment of diseases beyond Spinal Muscular Atrophy. CP3 defines a new class of compounds that activate weak 5’ ss (GU|GU) other than the GA|GU 5’ ss activated by currently available small molecule splicing modulators (risdiplam and branaplam). Mechanistically, CP3 and risdiplam-like molecules enhance U1 snRNP binding to weak 5’ ss through recruitment of Luc7 or Luc7L2, whereas branaplam acts independently of these cofactors.^40^ The identification of residues on Luc7L critical for the effect of CP3 support a conserved mechanism in which Luc7L family of proteins engage the U1 and 5′ ss duplex analogously to yeast Luc7, an integral component of yeast U1 snRNP, thereby revealing the molecular mechanism of this class of alternative splicing factors. The dependence of CP3 activity on Luc7L/L2 may increase their specificity to cells that express these alternative splicing factors and provide us with an opportunity to use Luc7L activators to potentiate these compounds.

Beyond prion disease, several additional disease-modifying genes, such as *LRRK2*, were also identified as responsive to CP3, highlighting the broader therapeutic potential of this approach (Extended Data Fig. 13). Together, our findings establish a generalizable framework that integrates motif-directed medicinal chemistry with combinations of splicing modulators of different mechanisms, offering a roadmap to target proteins previously considered “undruggable,” particularly intrinsically disordered and scaffolding proteins.

## Methods

### Cell culture and transfection

HEK293T cells (Thermo Fisher, R70007) were cultured in DMEM (Gibco, 11995040) supplemented with 10% FBS (Cytiva, SH30910.03) and 1% Antibiotic-Antimycotic (Gibco, 15240062) at 37 °C in a humidified atmosphere with 5% CO₂. LN229 cells (ATCC, CRL-2611) were maintained in DMEM supplemented with 5% FBS and 1% Antibiotic-Antimycotic at 37 °C in 5% CO₂. Neuro-2A cells (ATCC, CCL-131) were cultured in DMEM supplemented with 10% FBS and 1% Antibiotic-Antimycotic at 37 °C in 5% CO₂.

For plasmid transfection, cells were transfected using Lipofectamine 2000 (Thermo Scientific, 11668019) with 1 μg plasmid DNA and 4 μL Lipofectamine 2000 per well in a 6-well plate, according to the manufacturer’s protocol. For siRNA and small oligonucleotide transfection, Lipofectamine RNAiMAX (Thermo Scientific, 13778150) was used with 1 μg oligonucleotide and 5 μL RNAiMAX per well in a 6-well plate, according to the manufacturer’s instructions.

### Splicing reporter assays

For *SMN2* splicing with 5′ss variants, plasmids of *SMN2* splicing cassettes (mutated from Addgene, #218670) were transfected into HEK293T cells. 12–16 hours after transfection, reporter cells were seeded in 384-well plates with 10,000 cells per well in a total of 30 μL of full medium. To determine dose responses, the cells were treated with a serial dilution of compounds at 12 concentrations in quadruplicate for 48 h before 10 μL of 0.5 × QUANTI-Luc (InvivoGen, rep-qlc2) was added to each well. The plates were measured immediately using a luminescence plate reader. *PRNP* splicing reporter assay was performed similarly. For RNA and protein quantification, *PRNP* reporter cells were seeded in 12-well plates with 50–60% confluency in a total of 1 mL medium, before treated with compounds. 24 hours after transfection, cells were harvested in RLT Plus buffer (QIAGEN, 1053393) for RNA extraction. Alternatively, 24-48 hours after transfection as indicated, cells were lysed in 1× RIPA buffer (Thermo Scientific, 89901) for Western blotting analysis.

### Endpoint RT-PCR and RT-qPCR analysis for *PRNP* mRNA

Total RNA was extracted from cells using RNeasy Mini kit (Qiagen, 74104). First strand cDNA was synthesized from total RNA, oligo(dT) primer, and reverse transcriptase (Promega, M1705) according to the manufacturer’s protocol. For endpoint RT-PCR, cDNA samples were amplified using PCR reaction (10 µL; Thermo Scientific, F124L) with 1 µL of 5-fold diluted cDNA sample according to the manufacturer’s protocol. PCR products were separated on 8% native polyacrylamide gels. Primer sequences: PRNP-Fw: 5’-AGCTTCTCCTCTCCTCACGA; PRNP-Rv: 5’-GTGTTCCATCCTCCAGGCTTC.

For RT-qPCR, 1.5 μL of 5-fold diluted cDNA was used in a 15 μL qPCR reaction with SYBR Green master mix (Apex Bio, K1070), following the manufacturer’s instructions. Primer sequence: PRNP-Fw: 5’-ACCGAGGCAGAGCAGTCATTAT; PRNP-Rv: 5’-GTGTTCCATCCTCCAGGCTTC. GAPDH-Fw: 5’-GCATCCTGGGCTACACTGAG; GAPDH-Rv: 5’-AAGTGGTCGTTGAGGGCAAT.

### Western blotting

Cells or homogenized tissues were lysed in 1× RIPA buffer (Thermo Scientific, 89901) with protease inhibitor cocktail (Thermo Scientific, PIA32955). The lysate was vortexed for 30 s and centrifuged at 17,000 g at 4 °C for 15 min to remove debris. The supernatant was mixed with 4× LDS Sample Buffer (Thermo Scientific, PIA32955), 50 mM DTT, then heated at 70 °C for 10 min. Western blotting was performed using Bolt 4–12% Bis-Tris gels (Invitrogen), followed by protein transfer using the iBlot 2 Gel Transfer Device (Invitrogen) according to the manufacturer’s instructions. Membranes were blocked in PBST containing blocking reagent and incubated with primary antibodies overnight at 4 °C or for 1 h at room temperature with gentle rocking. After washing with PBST, membranes were incubated with HRP-conjugated secondary antibody (1:2,500) for 1 h at room temperature, followed by additional washes. Signals were developed using HRP substrate (Thermo Scientific, 34580) and detected by chemiluminescence. Primary antibodies used were anti-PrP (Sigma, MAB1562; 0.1 µg/mL) and anti-FLAG M2 (Sigma, F3165; 10 µg/mL). Secondary antibody used was anti-mouse IgG HRP conjugate (Cell Signaling Technology, 7076).

### RNA-seq analysis

RNA-seq reads were preprocessed with fastp and aligned to the human reference genome (GRCh38, Gencode v44) using STAR in two-pass mode. Gene expression was quantified with featureCounts, normalized to log₂-TMM-CPM using edgeR, and tested for differential expression with a quasi-likelihood negative binomial GLM. Splice junction usage was quantified and tested for differential splicing using Leafcutter. To characterize dose-response relationships from RNA-seq, gene expression log2 (fold changes) were modeled as a three-parameter log-logistic function of dose using Bayesian inference (PyMC, NUTS sampler), with treatment (e.g. Risdiplam, CP3) -specific potency (EC_50_) and a shared maximum effect size. Junction-level splicing was similarly modeled with a four-parameter log-logistic curve and a Beta-Binomial likelihood to account for read-count overdispersion. Treatment-specific EC_50_ differences were identified via a specificity test on posterior distributions, corrected for genome-wide potency differences between treatments. Full details of computational methods, including model parameterizations, priors, filtering criteria, are provided in the **Supplementary Information**.

### Animal procedures

All animal procedures were reviewed and approved by the Institutional Animal Care and Use Committee (IACUC) of the University of Chicago, ensuring that all experiments were conducted ethically and in accordance with the NIH Guide for the Care and Use of Laboratory Animals. For this study, we used the human *PRNP* transgene on a *Prnp*^−/−^ background mice (Tg26372). Mice were maintained in ventilated cages under a 12-h light/dark cycle (light on 6 AM to 6 PM) and allowed access to food and water throughout the duration of the experiments.

### Primary neuronal culture

Primary neurons were isolated from humanized *PRNP* neonatal mice (P1–P3) using the papain dissociation system (Worthington, LK003150) following the manufacturer’s protocol. Cells (0.5–1.0 × 10^6^ per well) were plated on poly-*L*-lysine–coated 12-well plates in DMEM supplemented with 10% FBS and 1% Antibiotic-Antimycotic, then switched after 2–4 h to B-27 Plus Neuronal Culture System (Thermo Scientific, A3653401), supplied with 0.5 mM L-glutamine (Thermo Scientific, A2916801), and Penicillin/Streptomycin (Thermo Scientific, 15140122). Cells were treated with compounds after 1–2 days in culture and then subjected to splicing analysis and western blotting.

### In vivo pharmacodynamics

Neonatal mice (P1–P3) were anesthetized on ice and received intracerebral injections of 2 μL saline or drug solution into the right cerebral cortex. After recovery and rewarming, pups were returned to their dam. At 24 h post-injection of the indicated doses, mice were euthanized, and the injected right hemisphere was collected and snap-frozen on dry ice. For downstream analyses, brain tissues were homogenized in RLT Plus buffer (QIAGEN, 1053393) with stainless steel beads (QIAGEN, 69989). Total RNA was extracted from homogenate using RNeasy Mini kit (Qiagen, 74104), and proteins were precipitated from the RNeasy spin column flow-through of the same samples using acetone, according to the manufacturer’s instructions. To assess the in-vivo drug effect, human ***PRNP*** mRNA splicing was analyzed by endpoint RT-PCR; and prion protein levels were measured by western blotting.

### Fluorescence polarization assays for U1 snRNP binding to 5’ ss oligo

5’-FAM labeled PRNP RNA oligo was synthesized and purified by Integrated DNA Technologies (Coralville, USA). We performed FP assays in binding buffer (20 mM HEPES [pH 7.5], 1 mM MgCl_2_, 200 mM KCl, 0.01% TritonX-100, 0.1%PEG3350, 2%DMSO) at room temperature using an Envision HTS Plate Reader system (Revvity). Each reaction system contains 3 nM of FAM-labeled RNA oligo, reconstituted U1 snRNP in 2-fold increasing concentrations, 800 nM U1C, and different small molecules in DMSO (10 mM). 10 μM Luc7L/L2/L3 protein was incubated in dark for 30 min at room temperature. The equilibrium dissociation constants (*K*_d_) of the binding components were calculated using the following single-site binding model formula in GraphPad Prism:

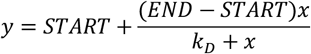

where x and y represent the concentration of U1 snRNP (or U1 snRNA) and the polarization value measured, respectively, START represents the polarization value of protein-free nucleic acid probe, END-START represents the maximum polarization value when U1 snRNP binds to the fluorescently labeled 5’ ss oligo. All FP experiments are carried out in two distinct biological replicates, and error bars represent the standard deviation.

## Acknowledgement

Research reported in this article was supported by the National Institutes of Health under award numbers R35GM147498 (J.W.), R35GM153249 (Y.I.L.), R35GM145289 (R.Z.), and R21NS142950 (R.Z.); the W. M. Keck Foundation (J.W., Y.I.L.); the CJD Foundation (J.W., J.A.M.); the Brain Research Foundation (J.A.M.); and the University of Chicago. Y.I.L. is a Biohub Investigator. Y.I.L. also thanks Sarah Yang and Frank Zeng for a research gift that helped support this work. We thank Sonia Vallabh and Eric Minikel (Broad Institute/Prion Alliance) for providing Tg26372 mice, Frédéric Allain (ETH Zurich) for providing U1 snRNP expression constructs, Frederic Vaillancourt and Bryan Dunyak at Remix Therapeutics, Inc. for providing Luc7 constructs and for useful discussions. We thank Xiaochang Zhang and Ming Xu (University of Chicago) for helpful discussions on animal experiments, and Xiangbin Ruan and Patrizia Lopresti (University of Chicago) for assistance with animal studies.

## Data Statement

All RNA-seq data generated in this study have been deposited in the NCBI Gene Expression Omnibus (GEO) database under accession number GSE330903. Processed data and associated metadata are available through the same repository.

---

**Extended Data Fig. 1:**
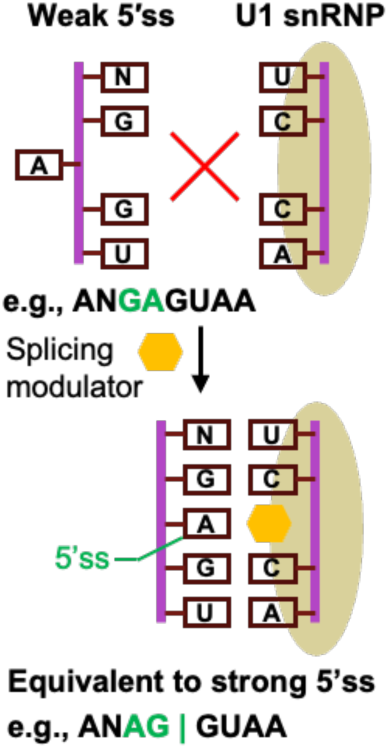
Molecular glue mechanism of splicing modulator that functions as an equivalent of splicing-favorable mutations on weak 5′ ss. As an illustration, a 5′ ss containing GA|GU is equivalent to AG|GU when treated with splicing modulators (e.g., risdiplam).

**Extended Data Fig. 2:**
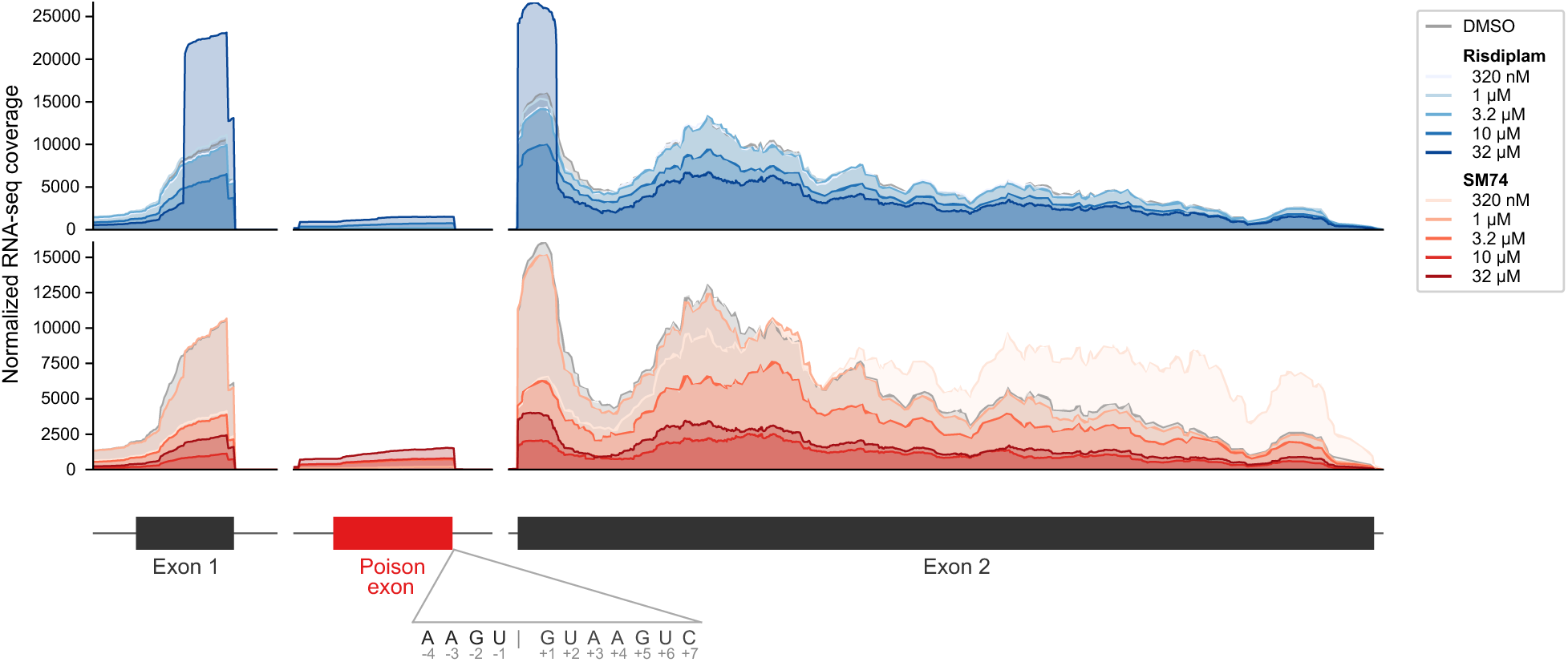
RNA-seq coverage tracks for *PRNP* cryptic exon inclusion induced by splicing modulators. RNA-seq read coverage across the *PRNP* gene in cells treated with various concentrations of risdiplam (blue) or SM74 (red). The poison exon is highlighted in red between exon 1 and exon 2. The sequence of the 5′ splice site of the cryptic exon is shown, with positions labeled relative to the splice junction.

**Extended Data Fig. 3:**
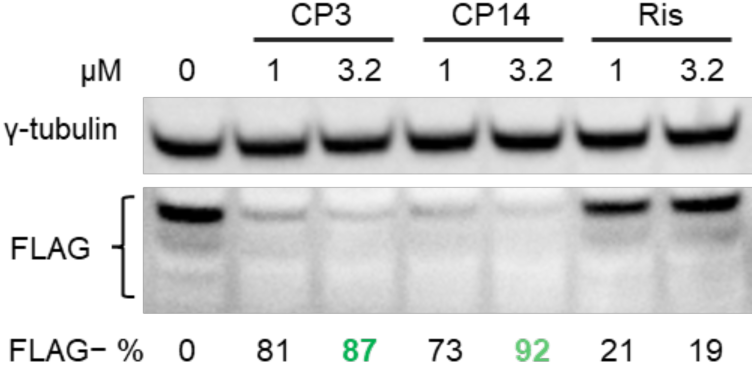
PrP expression suppression by anti-FLAG Western blot (24 h, PrP-FLAG minigene-transfected 293T cells, FLAG−%, percent reduction in PrP-FLAG levels). Data are representative of three independent experiments.

**Extended Data Fig. 4:**
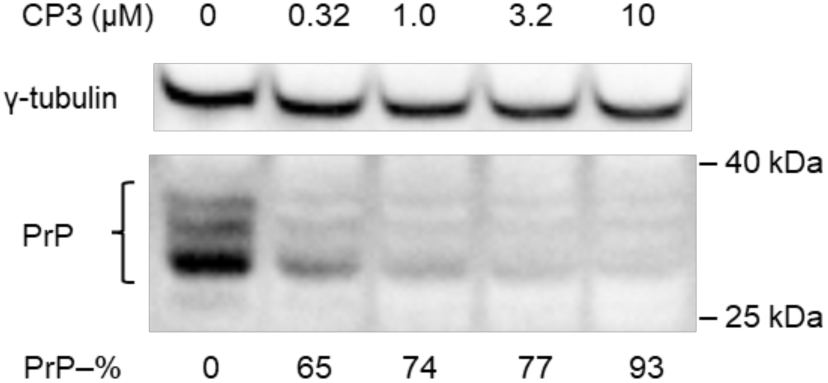
Dose-response activity of CP3 in Neuro-2A cells transfected with the PrP-FLAG minigene (24 h). PrP levels were measured by anti-PrP Western blot and quantified by densitometry. Data are representative of three independent experiments.

**Extended Data Fig. 5:**
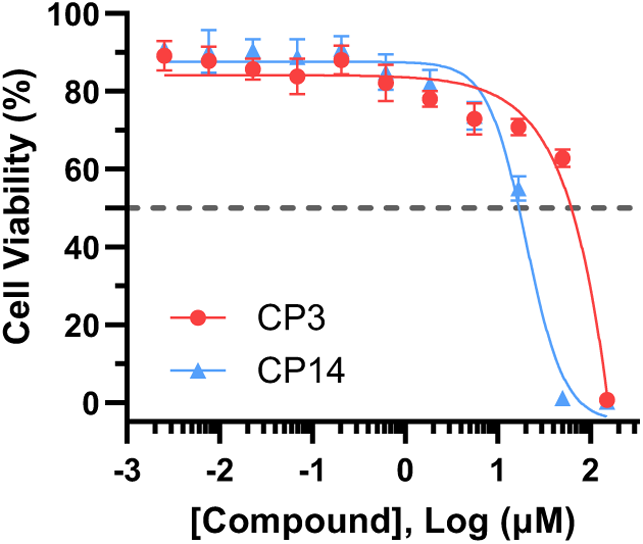
Cytotoxicity of CP3 and CP14 in 293T cells measured by CellTiter-Glo assay (48 h). Error bars represent standard deviation from three biological replicates.

**Extended Data Fig. 6:**
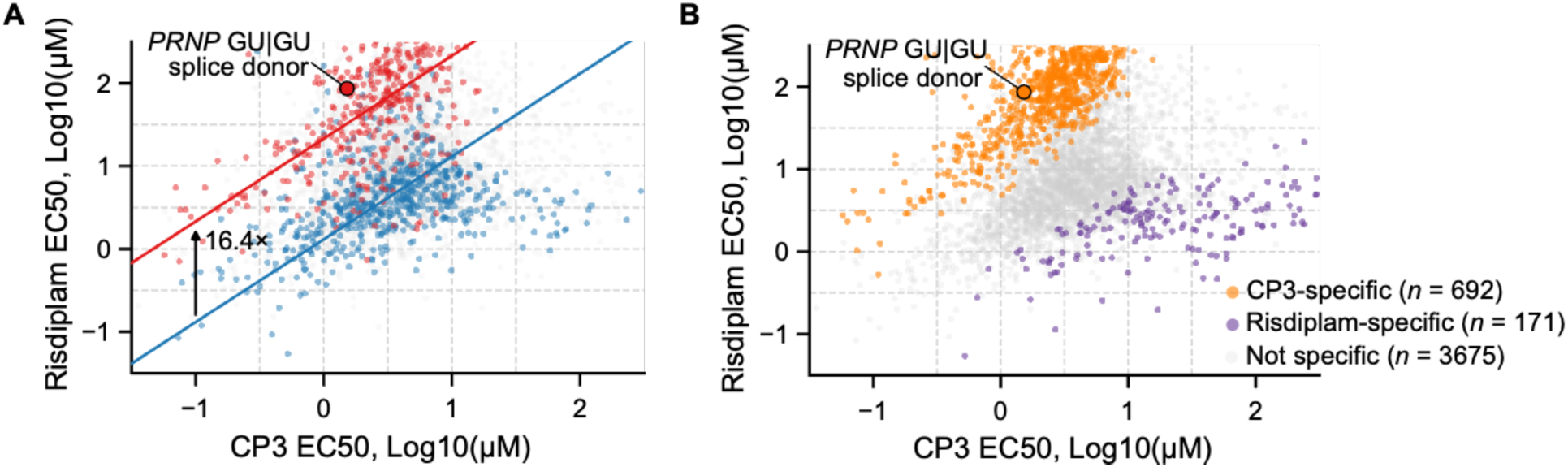
(A) Dose-response modeling reveals differences in 5′ ss activation between risdiplam and CP3. EC_50_ values were estimated for each differential splicing event (points colored red, blue, or gray for GA|GU, GU|GU, or other 5′ ss, respectively). RNA-seq assay doses are indicated by dashed lines. Median EC50 differences between CP3 and risdiplam were estimated for GA|GU (red trend line, linear regression) and GU|GU (blue trend line) splice junctions. (B) The scatter plot is recolored based on compound selectivity, defined by differences in EC_50_ between risdiplam and CP3 (local false sign rate < 0.01).

**Extended Data Fig. 7:**
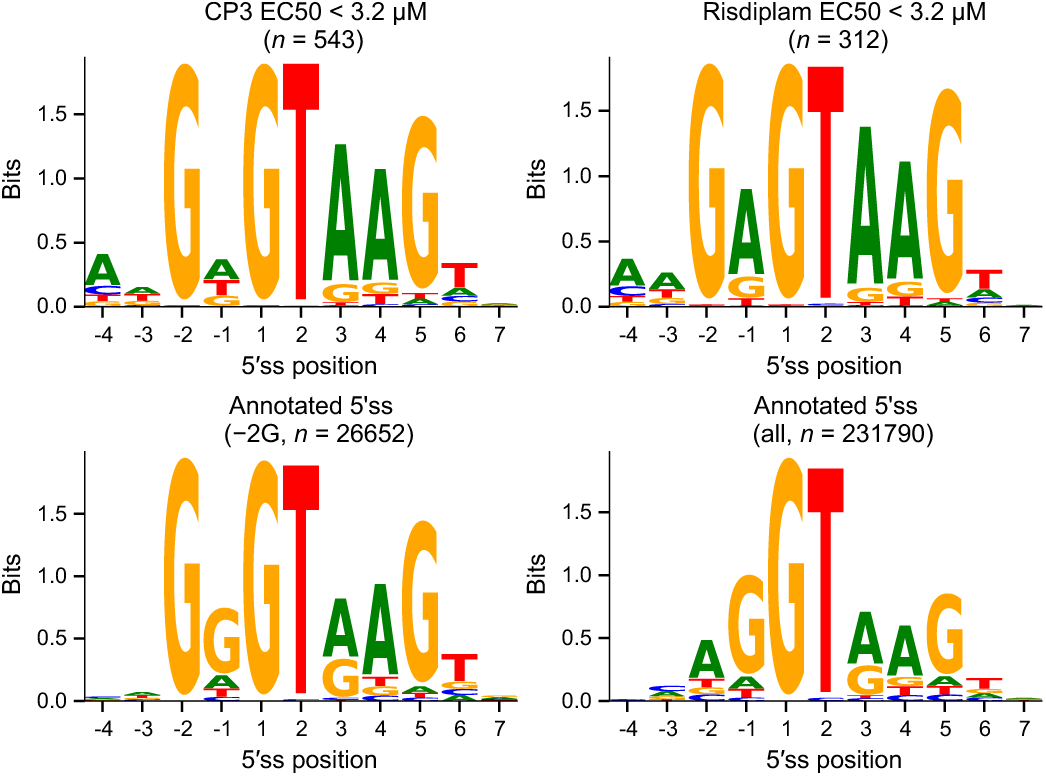
Seqlogos of 5′ ss from all CP3-activated (EC_50_ < 3.2 μM, *n* = 543) and risdiplam-activated (EC_50_ < 3.2 μM, *n* = 312) splice junctions, filtered for −2G to highlight direct splicing modulation (top). Seqlogos of all annotated 5′ ss (± −2G filtering) are shown for comparison (bottom).

**Extended Data Fig. 8:**
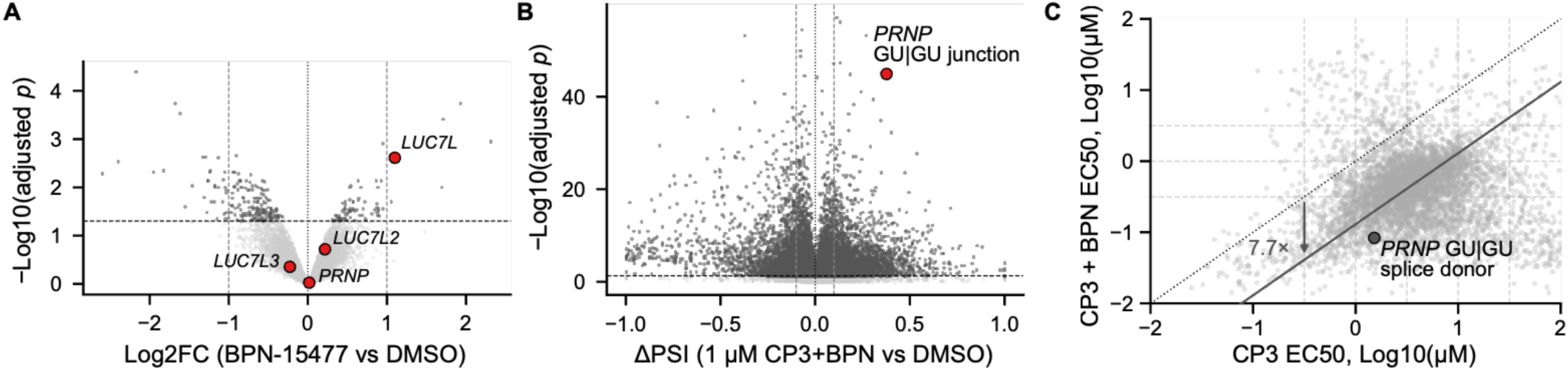
Volcano plots of (A) differential gene expression in cells treated with BPN-15477 (25 µM) versus DMSO control (24 h, LCL cells; dashed lines, adjusted *p*-value of 0.05 and Log2FC of ±1), and (B) differential splicing in cells treated with CP3 (1 μM) in combination with BPN-15477 (25 μM) versus DMSO control (24 h, LN229 cells; dashed lines, adjusted *p*-value of 0.01 and ΔPSI of ±10%. (C) Comparison of EC_50_ values derived from RNA-seq-based splicing analysis in LN229 cells treated with CP3 alone versus CP3 in combination with BPN-15477 (25 µM) across the genome. Each point represents an individual drug-responsive splicing event. The solid line represents a linear regression fit of all data, corresponding to an overall 7.7-fold increase in potency with the combination treatment (dashed line, *y = x*). The *PRNP* cryptic splicing with GU|GU splice donor is highlighted.

**Extended Data Fig. 9:**
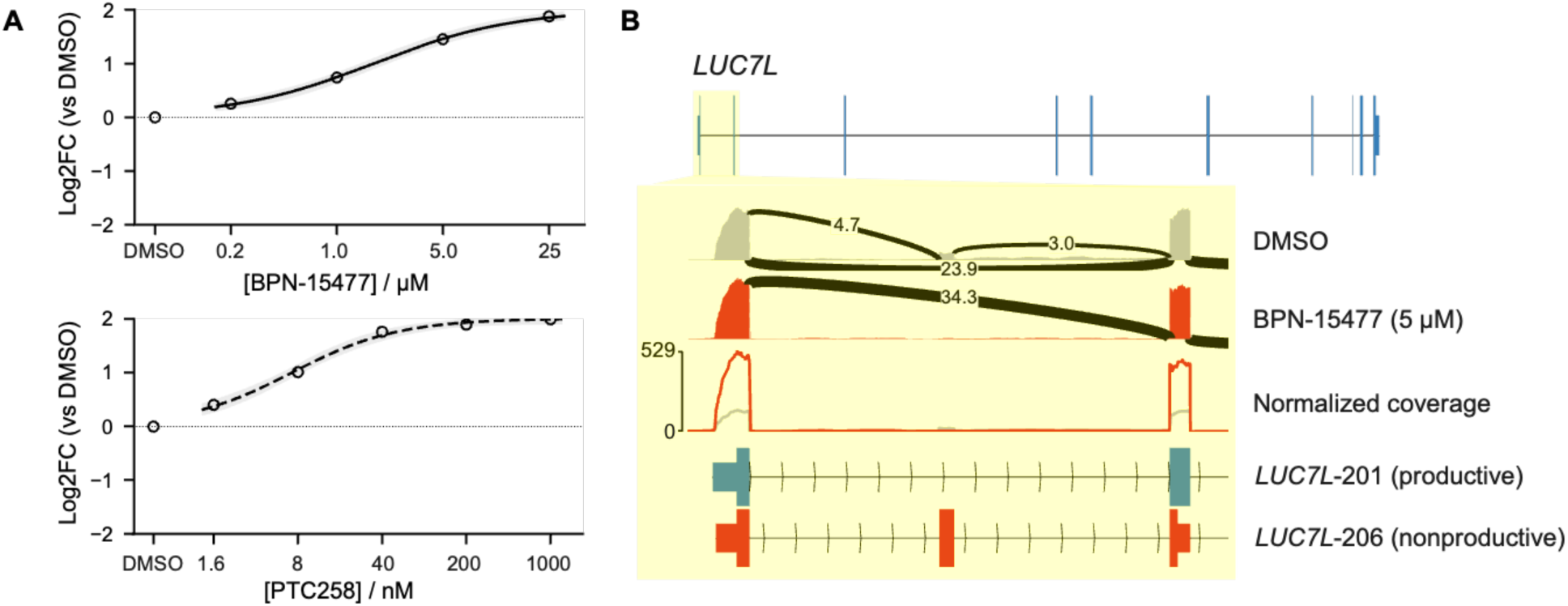
(A) *Luc7L* expression level changes estimated by RNA-seq analysis in LN229 cells treated with BPN-15477 (top) or PTC258 (bottom) for 24 h. Expression values were normalized to DMSO-treated controls. (B) Normalized base coverage per billion reads across the *Luc7L* gene reveals that intron 1 (yellow shade) contains a naturally occurring poison exon, resulting in >50% nonproductive transcripts (*LUC7L*-206 isoform). This mis-splicing event is corrected by BPN-15477 (5 µM, LCL cells) to restore the productive isoform (*LUC7L*-201). Sashimi plots are derived from the same coverage data but are auto-scaled for each track to enhance visualization of splice junction usage and the poison exon.

**Extended Data Fig. 10:**
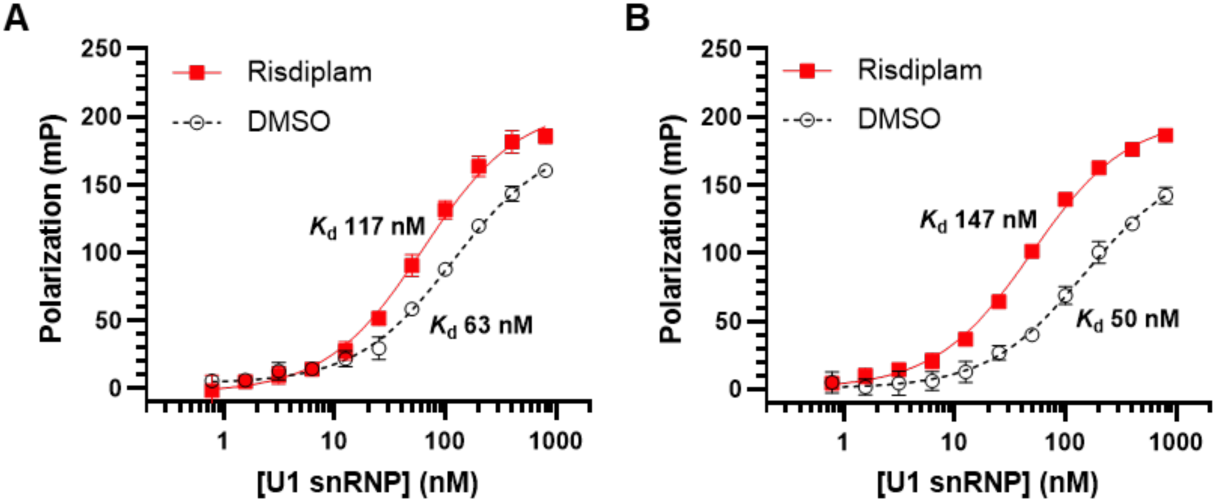
Luc7L and Luc7L2 enhance U1 snRNP−*SMN2* 5′ss−risdiplam complex formation. In vitro binding of U1 snRNP to *SMN2* pre-mRNA oligonucleotide (5’ 6-FAM-tagged) containing GA|GU 5′ss, measured by fluorescence polarization (FP) in the presence of risdiplam or DMSO control. Assays were performed with **(A)** Luc7L (10 μM) or **(B)** Luc7L2 (10 μM). Estimated *K*_d_ values from curve-fitting are shown beside each curve.

**Extended Data Fig. 11:**
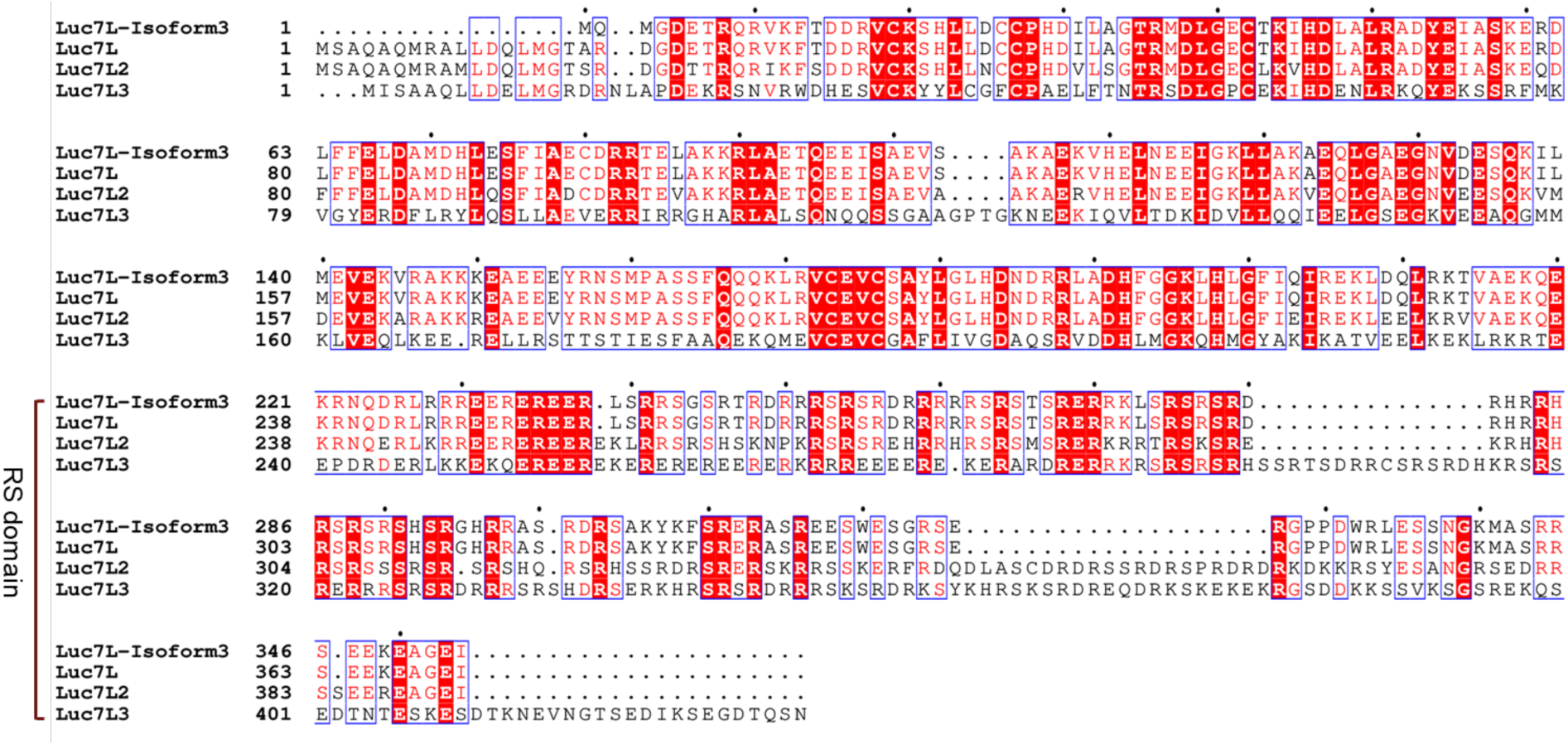
Multiple sequence alignment of Luc7L family proteins, including Luc7L isoform 1, Luc7L isoform 3, Luc7L2, and Luc7L3. Conserved residues are highlighted in red, with highly conserved positions indicated by boxed regions.

**Extended Data Fig. 12:**
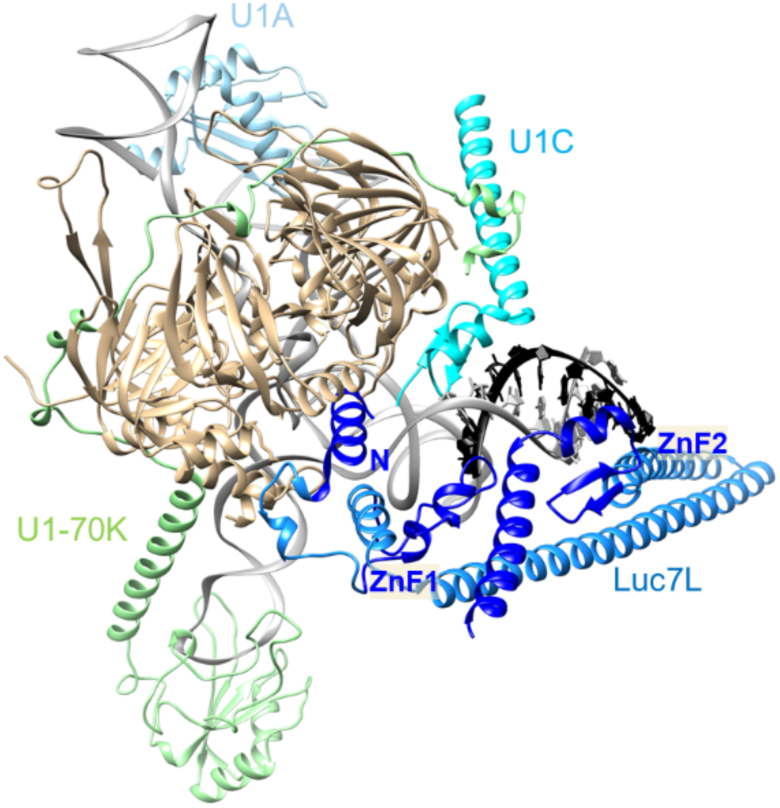
AlphaFold3 model indicates that Luc7L contacts the Sm ring through its N-terminal helix and the U1 and 5′ ss RNA duplex through ZnF2.

**Extended Data Fig. 13:**
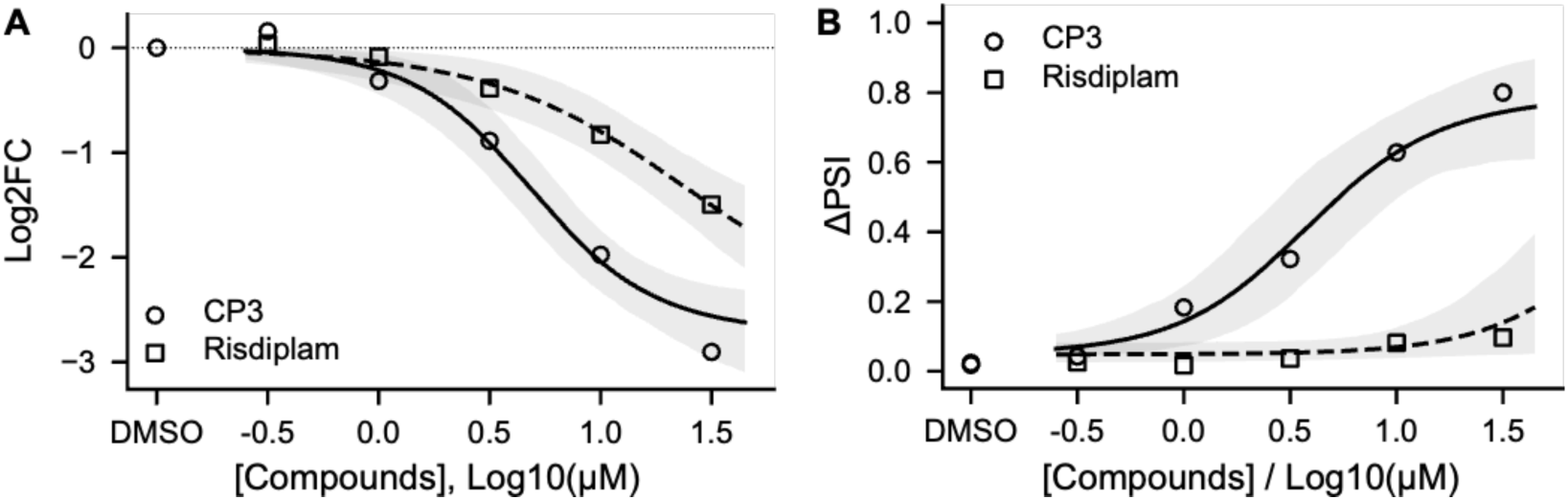
Dose response curves of CP3 and risdiplam for *LRRK2* in LN-229 cells (24 h treatment), showing (A) changes in expression level (log2 fold change) and (B) cryptic splicing (ΔPSI), derived from RNA-seq analysis.

